# Comparative analysis of single-cell RNA Sequencing Platforms and Methods

**DOI:** 10.1101/2020.07.20.212100

**Authors:** John M. Ashton, Hubert Rehrauer, Jason Myers, Jacqueline Myers, Michelle Zanche, Malene Balys, Jonathan Foox, Chistopher E. Mason, Robert Steen, Marcy Kuentzel, Catharine Aquino, Natàlia Garcia-Reyero, Sridar V. Chittur

## Abstract

Single-cell RNA sequencing (scRNA-seq) offers great new opportunities for increasing our understanding of complex biological processes. In particular, development of an accurate Human Cell Atlas is largely dependent on the rapidly advancing technologies and molecular chemistries employed in scRNA-seq. These advances have already allowed an increase in throughput for scRNA-seq from 96 to 80,000 cells on a single instrument run by capturing cells within nano-liter droplets. While this increase in throughput is critical for many experimental questions, a thorough comparison between microfluidic-, plate-, and droplet-based technologies or between multiple available platforms utilizing these technologies is largely lacking. Here, we report scRNA-seq data from SUM149PT cells treated with the histone deacetylase inhibitor TSA vs. untreated controls across several scRNA-seq platforms (Fluidigm C1, WaferGen iCell8, 10X Genomics Chromium Controller, and Illumina/BioRad ddSEQ). The primary goal of this project was to demonstrate RNA sequencing (RNA-seq) methods for profiling the ultra-low amounts of RNA present in individual cells, and this report discusses the results of the study as well as technical challenges/lesson learned and present general guidelines for best practices in sample preparation and analysis.

## INTRODUCTION

Cells are the most fundamental building blocks of all living organisms. From a unicellular bacteria to complex living forms composed of many different cell types, the transcriptional signatures of individual cells reflect the physiological as well as pathological state of a biological being. While analyses at the single-cell level have long been practiced in biology and medicine through microscopic observations, protein immunohistochemistry, *in situ* hybridization, and flow cytometry, the ability to perform whole transcriptome profiling (RNA-seq) by Next Generation Sequencing (NGS) at the resolution of single cells is a relatively recent and due to its ability to do parallel profiling of thousands of cells a very powerful approach. These new techniques enable novel discoveries and provide great insights into cell identity, function, cellular composition of different organs, as well as cell origin, evolution and heterogeneity in many different cancer types^1^.

In the past decade, different strategies have been explored for transcriptome profiling of single cells. Early on, single cell analyses by NGS were carried out using cell sorting to plate single cells into individual wells followed by cell lysis, cDNA synthesis, barcoded library generation, pooling and sequencing^2^. This method allows for individual cell assessment such as viability and individual cell capture prior to downstream processing. However, it is low throughput, labor intensive, and expensive. Utilizing integrated fluidic circuits, Fluidigm C1 autoprep system isolates single cells into individual nanochannels for visual examination followed by cell lysis, cDNA conversion, pre-amplification and retrieval for library construction and sequencing^3^. The C1 system significantly simplified the individual cell isolation while still enabling whole transcript sequencing analysis, however, cell partitioning is size restricted based on the nanochannel tolerance of the nanofluidic plate. DropSeq technology encapsulates single cells of different sizes in an oil-based droplet along with barcoded beads attached to unique oligomers in individual aqueous droplets for gene expression profiling of many hundreds to thousands of cells in parallel^4^. While DropSeq is highly efficient with commercially available platforms that are relatively easy to operate, only 5’- or 3’-tag profiling is possible at present, thereby not allowing for full-length transcript analysis. Furthermore, no intermediate assessment of cell capture quality and quantity is possible until after the completion of the NGS sequencing. The ICell8 platform, in contrast, isolates 1000-1800 cells in a 5184-nanowell chip to allow analysis of cell capture and viability^5^. Gene expression profiling by RNA-seq can be done for both 3’ profiling and full-length transcriptome. Currently, the most commonly employed microfluidics-based platform is the single-cell Chromium controller from 10x Genomics^6^. The 10x protocol is based on a 5’- or 3’-tag sequencing method. The Illumina/BioRad ddSEQ uses disposable microfluidic cartridges to co-encapsulate single cells and barcodes into subnanoliter droplets. Cell lysis and barcoding occur in the droplets, and libraries can then be subsequently prepared and sequenced

Considering each platform employs different strategies for single cell transcriptome profiling, the capacity, sensitivity, and reproducibility of each approach can differ greatly^7^. To better understand those strategies, the Association of Biomolecular Resource Facilities (ABRF) Genomics Research Group developed a study to compare different scRNA-seq platforms and methods. In order to reduce sample heterogeneity associated with dissociated cells from a given tissue type and to enable direct comparison of the platforms, we selected a single cancer cell line, SUM149 P/T, and tested gene expression signature changes after cells were treated with or without a HDAC inhibitor, trichostatin A (TSA) for this multi-platform comparison study^8^. We used bulk RNA-seq data as reference to assess the performance of five platforms including Fluidigm C1 and HT, 10X Chromium, BioRad ddSEQ, and ICELL8.

## MATERIALS AND METHODS

### Cell Culture and Drug Treatment

SUM149 P/T cells (gift from Dr. Martin Tenniswood, SUNY Albany) were grown in Ham’s F-12 medium (Invitrogen) supplemented with 5% FBS (Sigma-Aldrich), 5 µg/mL insulin (Sigma-Aldrich), 1 µg/mL hydrocortisone (Sigma-Aldrich), 10 mM HEPES (Fisher Scientific, Pittsburgh, PA) and antimycotic/antibiotic (Sigma-Aldrich).After the cells attached to the culture dishes, cells were grown and maintained in the above media with no FBS. Cells were maintained at 37 °C in a humidified atmosphere of 95% air/5% CO2. SUM149PT cells were plated at a density of 1.5 × 10^6^ cells per 150 cm^2^ dish for 48 h prior to treatment with 10nM TSA (in DMSO) or an equivalent volume of DMSO respectively. After 48 h of treatment, cells were harvested by trypsinization, washed with PBS and used for the scRNA-seq experiments.

### Fluidigm C1 96 scRNA-seq

Single SUM149 P/T cells treated with vehicle or TSA were captured on a 10–17μm cell diameter integrated microfluidic chip (IFC) using the Fluidigm C1 Single-cell AutoPrep system (Fluidigm Corporation). Cells were pre-stained with Calcien AM/EthD-1 LIVE/DEAD cell viability assay (Life Technologies) and loaded onto the IFC at a concentration of 500–700□cells/ul. Viable single cell confirmation was performed with phase-contrast fluorescence microscopy to assess the number and viability of cells per capture site. Only single, live cells were included in the analysis. For RNA-seq analysis, cDNAs were prepared “on-IFC” using the SMARTer Ultra Low RNA kit for Illumina (Clontech) following Fluidigm recommendations. Single-cell cDNA size distribution and concentration was assessed with PicoGreen (Life Technologies) and Agilent Bioanalyzer 2100 analysis (Agilent Technologies). Illumina libraries were constructed in 96-well plates using Illumina’s NexteraXT DNA Sample Preparation kit following the protocol supplied by Fluidigm. For each C1 experiment, a bulk RNA control and a negative control were processed in parallel, using the same reagent mixes as used on chip. Libraries were quantified by Agilent Bioanalyzer, using High Sensitivity DNA analysis kit, and also fluorometrically, using Qubit dsDNA HS Assay kits and a Qubit 2.0 Fluorometer (Life Technologies). Single end reads of 100nt were generated for each sample using Illumina’s HiSeq2500v4.

### Fluidigm C1 HT scRNA-seq

Single SUM149 P/T cells treated with vehicle or TSA were captured on a HT 10–17μm cell diameter integrated microfluidic chip (IFC) using the Fluidigm C1 Single-cell AutoPrep system (Fluidigm Corporation). Cells were pre-stained with Calcien AM/EthD-1 LIVE/DEAD cell viability assay (Life Technologies) and loaded onto the IFC at a concentration of 400□cells/ul. Viable single cell confirmation was performed with phase-contrast fluorescence microscopy to assess the number and viability of cells across 10% of the capture wells. For RNA-seq analysis, cDNAs were prepared “on-IFC” using the SMARTer Ultra Low RNA kit for Illumina (Clontech) following Fluidigm recommendations. Single-cell cDNA size distribution and concentration was assessed with PicoGreen (Life Technologies) and Agilent Bioanalyzer 2100 analysis (Agilent Technologies). Illumina libraries were constructed in 96-well plates using Illumina’s NexteraXT DNA Sample Preparation kit following the protocol supplied by Fluidigm. For each C1 experiment, a bulk RNA control and a negative control were processed in parallel, using the same reagent mixes as used on chip. Libraries were quantified by Agilent Bioanalyzer, using High Sensitivity DNA analysis kit, and also fluorometrically, using Qubit dsDNA HS Assay kits and a Qubit 2.0 Fluorometer (Life Technologies). Single end reads of 100nt were generated for each sample using Illumina’s HiSeq2500v4.

### WaferGen iCell8 scRNA-seq

Cells were stained with Hoechst 33324 and Propidium Iodide (Thermo Fisher Scientific) for 20 min. The cell viability and density was checked with Moxi Cell counter (VWR) and cells were diluted to achieve a density of 1 cell per 50 nl in a final dispensing mix which contained a diluent, murine RNase inhibitor (New England Biolabs) and 0.35X PBS (without Ca^++^ and Mg^++^, pH 7.4, Thermo Fisher Scientific). A 384-well source plate with 8 designated wells containing cell suspensions, 1 well for positive control, 1 well for negative control, and 1 well for fiducial mix (fluorescent dye permitting image alignment confirmation) was placed in the Multisample nanodispenser (MSND, WaferGen). Each of the 8 sample source wells in the 384-source plate was sampled by 1 of the 8 channels in MSND. Cells, positive controls, negative controls, and fiducial mix, were dispensed onto one chip within 16 min. Each well received 50 nl of either cell mix, positive control, negative control, or fiducial mix. Two ICELL8 chips were used to collect single cells after which the chips were sealed, centrifuged at 300g for 5 min at room temp. The chips were subsequently imaged using the imaging station and frozen at -80°C overnight. Microchip images were analyzed using CellSelect software (WaferGen) to determine the viability and number of cells present in each nanowell. From each chip we selected 425 TSA treated single cells, 425 DMSO treated single cells, 4 positive controls and 4 negative controls.

Microchips were removed from -80 °C and left at room temperature for 10 min. Cells were lysed by freeze-thaw at this step. Chips were then centrifuged at 3800 g for 5 min at 4 °C and transferred to a thermocycler with a program of 72 °C for 3 min and 4 °C hold to anneal preprinted oligonucleotides to polyA mRNAs. The microchips were centrifuged as previously and were placed back into the MSND. A separate 384-well source plate containing reverse transcription (RT) reagents (GC Melt (5 M), dNTP Mix (25 mM each), MgCl2 (1 M), DTT (100 mM), 5X First-Strand Buffer, Triton X-100 (10%), SMARTer ICELL8 3’ DE Oligo Mix, SMARTScribe Reverse Transcriptase (100 U/μl), SeqAmp DNA Polymerase) in 4 wells was used in the MSND, which delivered 50 nl of reverse transcription mix to selected nanowells. The microchips were spun down and transferred to a Chip cycler to perform RT-PCR using the preinstalled program (500C for 3 min, 4oC for 5 min, 42 °C for 90 min, 2 cycles of 50ºC 2min and 42°C 2 min followed by heating at 70°C for 15 min, 95°C for 1min, 24 cycles of 98°C 10sec, 65°C 30 sec, 68°C 3 min and finally 72°C for 10 min followed by ramp down to a 4 °C hold. Post reaction chips were inverted and centrifuged (3800 g 10 min at 4 °C) to simultaneously collect and pool well contents into a single microcentrifuge collection tube. Double-stranded cDNA was cleaned by the DNA Clean & Concentrator™-5 kit (Zymo Research) purified using a 0.6X proportion of AMPure XP beads. (Beckman Coulter). cDNA quality was assessed using a Bioanalyzer High Sensitivity DNA chip (Agilent Technologies) and quantity was determined by a Qubit High Sensitivity kit (Thermo Fisher Scientific). Sequencing libraries were made using the Nextera XT DNA Library Preparation Kit (Illumina) that added Illumina adapters and indexes to the purified cDNA via a tagmentation reaction followed by PCR. The sequenced ready libraries were checked for quality and quantity using an Agilent High sensitivity BioAnalyzer assay, a dsDNA HS qubit assay and also by a NEBNext library quantitation Assay (New England Biolabs) before sequencing on Illumina’s HiSeq2500v4

### 10X Genomics scRNA-seq

Cellular suspensions were loaded on a Chromium Single-Cell Instrument (10x Genomics, Pleasanton, CA, USA) to generate single-cell Gel Bead-in-Emulsions (GEMs). Single-cell RNA-seq libraries were prepared using Chromium Single-Cell 3′ Library & Gel Bead Kit (10x Genomics). The beads were dissolved and cells were lysed per manufacturer’s recommendations. GEM reverse transcription (GEM-RT) was performed to produce a barcoded, full-length cDNA from poly-adenylated mRNA. After incubation, GEMs were broken and the pooled post-GEM-RT reaction mixtures were recovered and cDNA was purified with silane magnetic beads (DynaBeads MyOne Silane Beads, PN37002D, Thermo Fisher Scientific). The entire purified post GEM-RT product was amplified by PCR. This amplification reaction generated sufficient material to construct a 3’ cDNA library. Enzymatic fragmentation and size selection was used to optimize the cDNA amplicon size and indexed sequencing libraries were constructed by End Repair, A-tailing, Adaptor Ligation, and PCR. Final libraries contain the P5 and P7 priming sites used in Illumina bridge amplification. Sequence data was generated using Illumina’s HiSeq2500v4.

### Illumina SureCell/ddSEQ scRNA-seq

Single SUM149 P/T cells treated with vehicle or TSA were filtered with 22-μm filter (Millipore) and kept in cold PBS-BSA solution. Cell viability was determined by trypan blue staining and analyzed using a BioRad TC20 Automated Cell Counter. Cell suspensions were loaded on the ddSEQ droplet generator at a final dilution of 2,500 cells/uL and containing no less than 90% live cells. Illumina SureCell WTA reagents for encapsulation, cDNA synthesis, and library construction were used. Cell suspensions were loaded into a ddSEQ cartridge in a ratio of 21.5uL cell enzyme mix to every 4.5uL cell suspension. A total of 60ul of 3′ barcode suspension mix and encapsulation oil were also loaded on the cartridge. The chip was then processed on a ddSEQ Single-Cell Isolator to generate single cell droplets. Droplet encapsulated cells were transferred to a 96-well plate and reverse transcription mix was added and followed by reverse transcription on a thermal cycler. Post reverse transcription, droplets were broken to release cDNA and followed by the first strand purification using magnetic beads. The beads were immobilized using magnetic peg stand and dynamag and supernatant was discarded, beads washed twice with 80% EtOH, cDNA eluted with resuspension buffer, and transferred to a fresh microfuge tube. The purified first strand was used to synthesize the second strand to generate the double strand cDNA. Amplified cDNA was simultaneously fragmented and barcoded by tagmentation using Nextera Tn5 on thermal cycler. Illumina DNA adapters were added, followed by the indexing. Final libraries were cleaned up using AmpureXP beads. Purified libraries were eluted with 20 ul resuspension buffer. Final libraries were assessed for quality and quantity with Qubit 2.0 flourometer and an Agilent BioAnalyzer 2100. Libraries were sequenced using an Illumina HiSeq 2500v4 sequencer using SureCell sequencing primers.

### TruSeq Bulk RNA scRNA-seq

The total RNA concentration was determined with the NanopDrop 1000 spectrophotometer (NanoDrop, Wilmington, DE) and RNA quality assessed with the Agilent Bioanalyzer (Agilent, Santa Clara, CA). The TruSeq Stranded mRNA Sample Preparation Kit (Illumina, San Diego, CA) was used for next generation sequencing library construction per manufacturer’s protocols. Briefly, mRNA was purified from 200ng total RNA with oligo-dT magnetic beads and fragmented. First-strand cDNA synthesis was performed with random hexamer priming followed by second-strand cDNA synthesis using dUTP incorporation for strand marking. End repair and 3’ adenylation was then performed on the double stranded cDNA. Illumina adaptors were ligated to both ends of the cDNA, purified by gel electrophoresis and amplified with PCR primers specific to the adaptor sequences to generate cDNA amplicons of approximately 200-500bp in size. The amplified libraries were hybridized to the Illumina single end flow cell and amplified using the cBot (Illumina, San Diego, CA). Single end reads of 100nt were generated for each sample using Illumina’s HiSeq2500v4.

### Low Input RNA scRNA-seq

Total RNA was isolated using the RNeasy Plus Micro Kit (Qiagen, Valencia, CA) or Arcturus Pico Pure kit (Life Technnologies, Carlsbad, CA) per manufacturer’s recommendations. RNA concentration was determined with the NanopDrop 1000 spectrophotometer (NanoDrop, Wilmington, DE) and RNA quality assessed with the Agilent Bioanalyzer 2100 (Agilent, Santa Clara, CA). 1ng of total RNA was pre-amplified with the SMARTer Ultra Low Input kit v4 (Clontech, Mountain View, CA) per manufacturer’s recommendations. The quantity and quality of the subsequent cDNA was determined using the Qubit Flourometer (Life Technnologies, Carlsbad, CA) and the Agilent Bioanalyzer 2100 (Agilent, Santa Clara, CA). 150pg of cDNA was used to generate Illumina compatible sequencing libraries with the NexteraXT library preparation kit (Illumina, San Diego, CA) per manufacturer’s protocols. The amplified libraries were hybridized to the Illumina single end flow cell and amplified using the cBot (Illumina, San Diego, CA). Single end reads of 100nt were generated for each sample using Illumina’s HiSeq2500v4.

### scRNA-seq data processing

All single-cell data was processed per vendor recommended or provided software tools to assess performance under ideal conditions:

#### ddSEQ/SureCell WTA

Data processing was performed with Illumina SureSelect WTA BaseSpace application. Briefly, raw data were demultiplexed using bcl2fastq version 1.8.4. Quality filtering and adapter removal were performed using Trimmomatic version 0.36 with the following parameters: “TRAILING:13 LEADING:13 ILLUMINACLIP:adapters.fasta:2:30:10 SLIDINGWINDOW:4:20 MINLEN:15”. Cell debarcoding and unique molecular identifiers (UMIs) identification was performed with the SureCell BaseSpace app using vendor recommended parameters. Reads that had missing pairs and cells with barcodes that could not be assigned to known barcode combinations were removed from downstream analysis. Reads assigned to each cell were aligned to Genome Reference Consortium Human Build 38 (GRCh38.7) library (Gencode-25) using Spliced Transcript Alignment to a Reference (STAR) algorithm (v2.5.2b) and duplicated reads with the same UMI aligning to the same genomic position were filtered to minimize amplification bias. Gene annotation was performed using featureCounts. Gene expression levels were calculated by counting the number of distinct UMIs of all gene specific reads (‘UMIs per gene’), as determined by featureCounts.

#### Fluidigm C1 and HT

Raw data were demultiplexed using bcl2fastq version 1.8.4. Quality filtering and adapter removal were performed using Trimmomatic version 0.36 with the following parameters: “TRAILING:13 LEADING:13 ILLUMINACLIP:adapters.fasta:2:30:10 SLIDINGWINDOW:4:20 MINLEN:15”. Sequencing data were cleaned using Trimmomatic, and aligned to Genome Reference Consortium Human Build 38 (GRCh38.7) library (Gencode-25) using Spliced Transcript Alignment to a Reference (STAR) algorithm (v2.5.2b). Uniquely aligned reads were quantified and quality-assessed using picardtools (v1.114). Outlier cells were removed based on the number of genes expressed (<2,000 or >10,000), percent mitochondrial (0–0.2) or ribosomal (0–0.1) reads. Gene annotation was performed using featureCounts.

#### WaferGen iCell8

Raw data were demultiplexed using bcl2fastq version 1.8.4. Quality filtering and adapter removal were performed using Trimmomatic version 0.36 with the following parameters:”TRAILING:13 LEADING:13 ILLUMINACLIP: adapters.fasta:2:30:10 SLIDINGWINDOW:4:20 MINLEN:15”. Sequencing data were cleaned using Trimmomatic, and aligned to Genome Reference Consortium Human Build 38 (GRCh38.7) library (Gencode-25) using Spliced Transcript Alignment to a Reference (STAR) algorithm (v2.5.2b). Uniquely aligned reads were quantified and quality-assessed using picardtools (v1.114). Outlier cells were removed based on the number of genes expressed (<2,000 or >10,000), percent mitochondrial (0–0.2) or ribosomal (0–0.1) reads. Gene annotation was performed using featureCounts. Gene expression levels were calculated by counting the number of distinct UMIs of all gene specific reads (‘UMIs per gene’), as determined by featureCounts.

#### 10X Genomics

Raw data were demultiplexed using bcl2fastq version 1.8.4. Quality filtering and adapter removal were performed using Trimmomatic version 0.36 with the following parameters: “TRAILING:13 LEADING:13 ILLUMINACLIP:adapters.fasta:2:30:10 SLIDINGWINDOW:4:20 MINLEN:15”. Quality filtered data Samples are demultiplexed using 10X Genomics recommended workflow via cellRanger mkfastq. The data were then aligned and counted using cellRanger count.

### Bulk RNA-seq data processing

Raw reads generated from the Illumina HiSeq2500 sequencer were demultiplexed using bcl2fastq version 2.19.0. Quality filtering and adapter removal are performed using Trimmomatic-0.36 with the following parameters: “TRAILING:13 LEADING:13 ILLUMINACLIP:adapters.fasta:2:30:10 SLIDINGWINDOW:4:20 MINLEN:35” Processed/cleaned reads were then mapped to the human reference sequence (GRCh38.7) with STAR-2.6.0c with the following parameters: “--twopassMode Basic --runMode alignReads --genomeDir ${GENOME} --readFilesIn ${SAMPLE} --outSAMtype BAM SortedByCoordinate --outSAMstrandField intronMotif --outFilterIntronMotifs RemoveNoncanonical”. The subread-1.6.1 package (featureCounts) was used to derive gene counts given the following parameters: “-s 2 -t exon -g gene_name”. Differential expression analysis and data normalization was performed using DESeq2-1.16.1 with an adjusted p-value threshold of 0.05 within an R-3.4.1 environment.

### Expression analysis

All analyses were performed using R/Bioconductor. We used the package SingleCellExperiment to load and represent the scRNA-seq data as well as the bulk RNA-seq Data. Computation of quality metrics was performed using the *scater* package. We used the hard thresholds of 3 reads per gene to call a gene detected and we required 500 detected genes per cell to accept a cell. Additionally, we required at least 20000 reads on protein coding genes per cell. For the computation of GC content, we stratified the genes according to their GC content and defined genes with a GC content below 0.4 as *low-GC* and genes with a GC content above 0.6 as *high-GC*. As reference we used the genes with a GC content in the range 0.5–0.55. We then computed the fraction of detected genes in the *low-GC* group and compared it to the fraction of detected genes in the reference group. The GC bias plots show the log-ratio of the two fractions per cell. For the length bias, we applied the same strategy by using the thresholds <800nt for short genes, >2500 for long genes, and 1200-1800nt for the reference gene sets.

Correlation of expression values was computed as Spearman’s rank correlation. We used the edgeR package to compute differential expression. Thereby we made use of the TMM normalization to estimate the between sample normalization factors. Highly Variable Genes were computed using the Seurat package.

### Availability

Data generated by all technologies have been uploaded to GEO and are available under the accession GSE142652.

## RESULTS

### Generation of scRNA-seq Data Sets

Much of the variation observed in single-cell RNA Sequencing (scRNA-seq) gene expression data is due technical variation of the various methods utilized and basic biological variability^9, 10^. In order to reduce biological variation, we used a well establish triple negative breast cancer cell line (SUM149 P/T) to limit cell heterogeneity confounding cross platform results. A well-defined and robust gene signature of SUM149 P/T exposed to Trichostatin A (HDAC inhibitor) is already published^8^. We treated SUM149 P/T cells with vehicle (DMSO) or Trichostatin A (TSA) for 16 hours prior to harvesting cells for scRNA-seq across the various platforms tested. Importantly, bulk total RNA was sampled from each experimental batch of cells across platforms to evaluate both robustness and reproducibility of the treatment and to control for technical treatment batch effects. For all five platforms tested, duplicate cell captures were performed to control for and assess intra-platform technical variability (Figure 1). Additionally, all SUM149 P/T cells utilized in this study were derived from the same stock cell culture passage using a defined standard operating procedure to reduce technical batch effects from cell culture. This controlled experimental study was designed with intent to minimize as much variation as possible in order to most accurately assess scRNA-seq technologies.

**Figure 1:**
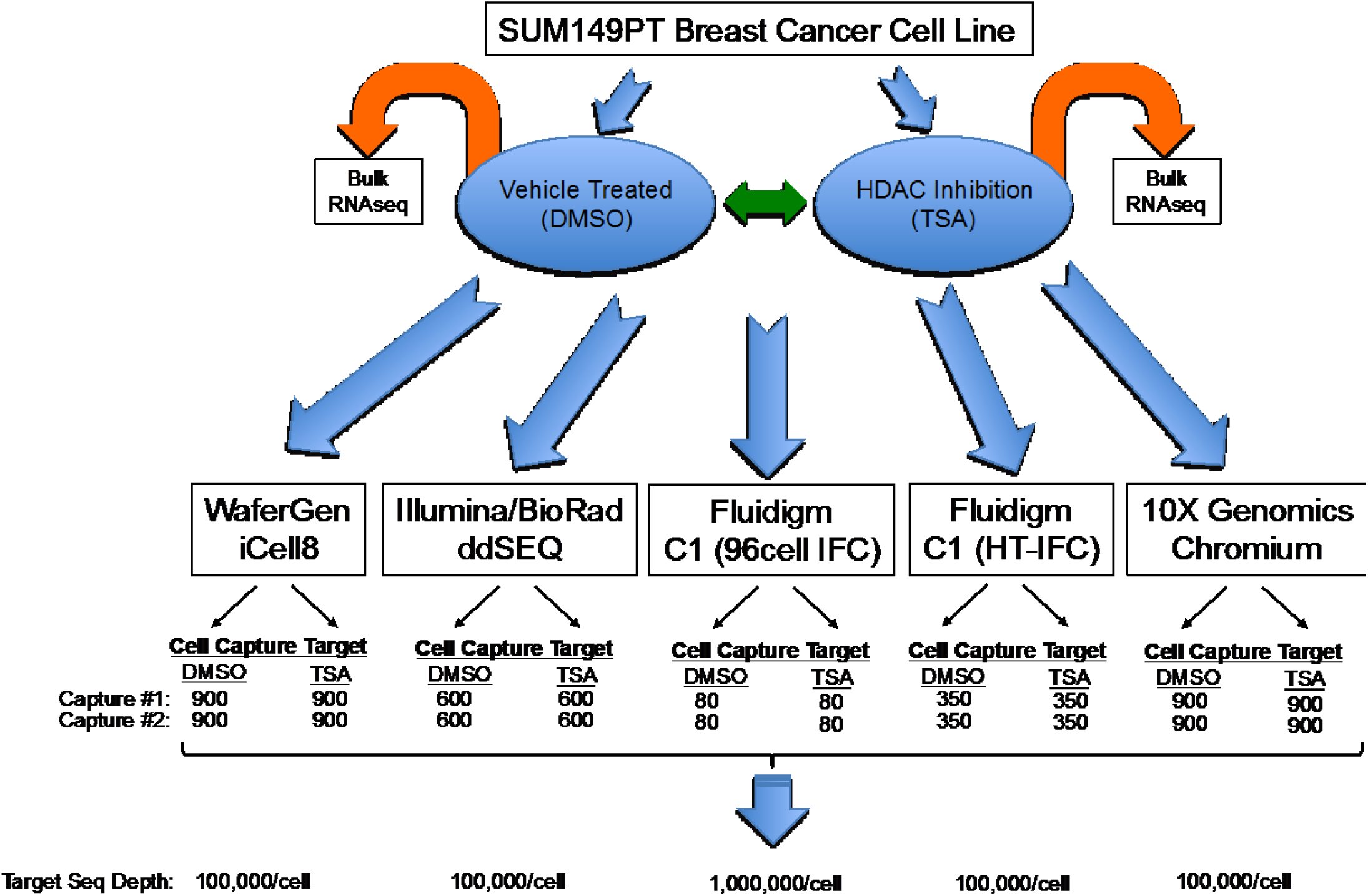
Experimental setup. Cells from the sum149PT breast cancer cell line were subjected to DMSO and TSA treatment and subsequently sequenced using different technologies.

### Throughput and sensitivity across platforms

While most platforms are capable of transcriptome-wide analysis, they vary in single-cell throughput from as little as 96 upwards of 80,000 single-cells. While each vendor claims a specific cell throughput, this is theoretical and influenced by many factors, such as cell viability, cell type, quality of cell suspension (i.e. presence of cellular debris). Given this, we sought to evaluate both cell capture throughput and gene detection sensitivity across platforms. In order to be as fair as possible to each technology, an attempt was made to keep read depth (sequence reads) per cell at vendor recommendations across technologies (Figure 2A). We next determined the number of unique genes detectable for each platform tested at the recommended sequencing depth (Figure 2B). Not surprisingly, technologies differ in the number of cells that can be processed in a single experiment (throughput) as well as in number of genes detectable, but interestingly, they also differ in the amount of “usable” data in terms of cells that pass quality control metrics and in terms of reads that actually do contribute to gene expression counts (see Figure 2C). Successful cell assessment can be impacted at the initial cell capture step, due to the stochastic nature of cell capturing between the different platform-specific protocols leading to absence of data those events. In addition to inefficient cell capture, each platform protocol may fail to generate sufficient reads from protein-coding genes necessary to generate reliable gene expression profiles. Given these potential issues, we used a fixed absolute threshold to assess the suitability of an expression profile. Specifically, we required a minimum of 20,000 reads per cell, greater than 500 genes detected per cell and a gene was counted as detected if at least 3 reads were assigned to that specific gene locus. Implementation of these absolute thresholds, imposed a minimum information requirement per cell for each technology that allowed a fairer assessment of each platform through use of only highly stringent cell and gene criteria. In summary, these observations suggest that some platform specific bias is present and that careful consideration should be given to this during experimental design to limit such confounding factors.

**Figure 2:**
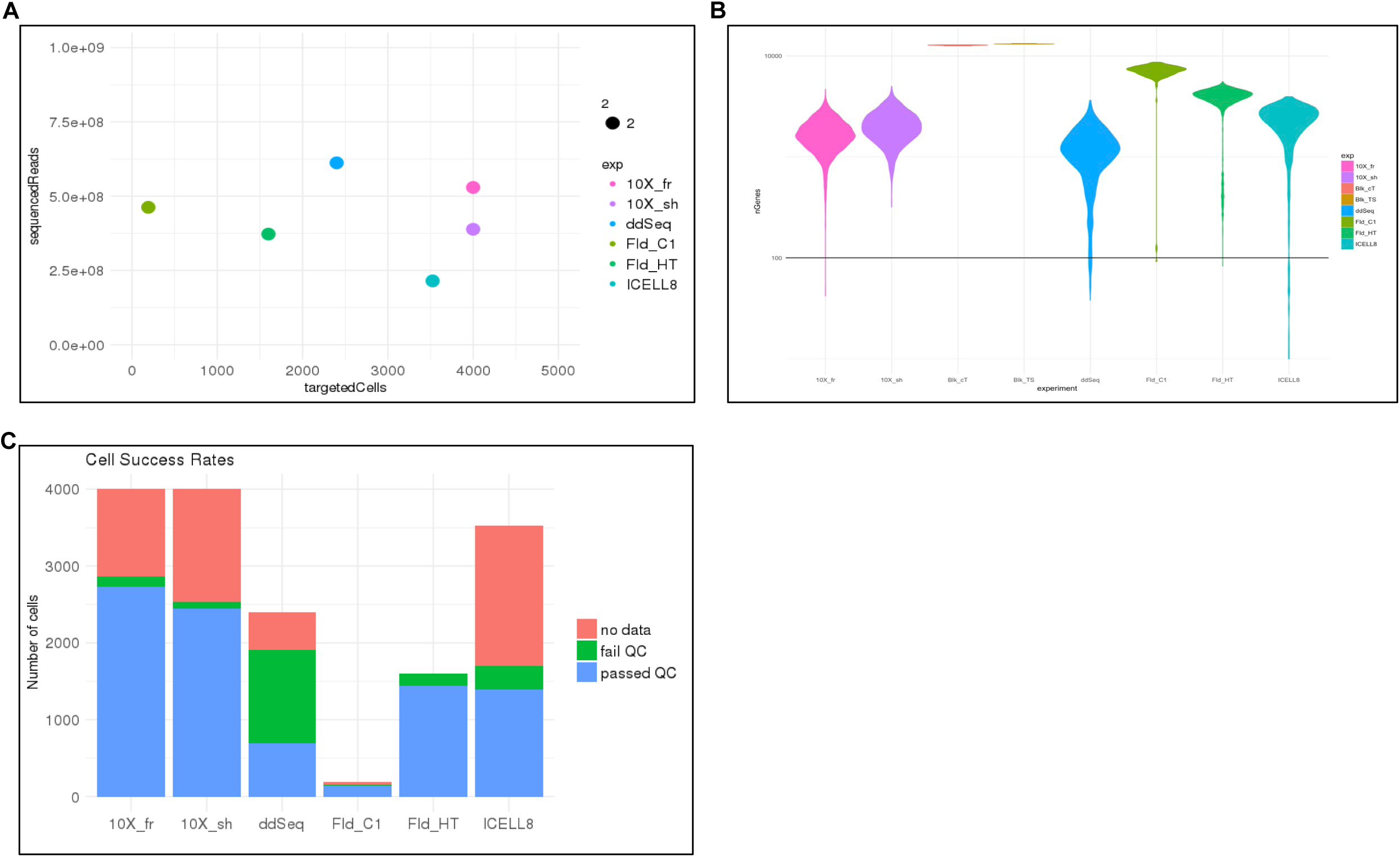
We show basic statistics of the single cell data. A: The plot shows for each technology the number of reads sequenced versus the number of cells targeted. 10X Genomics targeted the most cells. B: Violin plots showing the number of genes detected in all cells. The plot shows all cells without quality filtering. C: The bar plot show for each technology how if the targeted cells were actually detected and passed the QC filtering.

The total number of genes detected depends on the number of expressed genes and the transcript abundance in each cell type, as well as the sensitivity of the RNA capture and library preparation, and finally the sequencing depth. We obtained estimates of relative sensitivity of the different technologies by relating the number of genes detected in a cell to the number of UMIs/Reads sequenced for that cell. This provides information on the gene detection sensitivity as well as the saturation behavior. Our experiments show that, for the given cell type and respective sequencing depth, saturation was not yet reached for any of the platform technologies (Figure 3A-B). These data indicate that the number of detected genes per read depth, aligned and counted is similar for the different technologies. As shown in Figure 2B, the Fluidigm C1 methodology outperforms the other technologies at the recommended sequencing depth while ddSEQ has the lowest number of detected genes. However, these numbers do depend on the sequencing depth, which was chosen according to what is typically recommended for each scRNA-seq technology. As an alternative comparison, we show in Figure 3C the number of genes detected when considering only 20,000 reads per cell that were randomly sampled. Using the down-sampled dataset, all platforms performed comparably well with gene detection rates around 1,500, except for 10X that detected less than 1000 genes. These results suggest that in our study the diversity of the detected reads is lower for the 10X platform compared to the other platforms.

**Figure 3:**
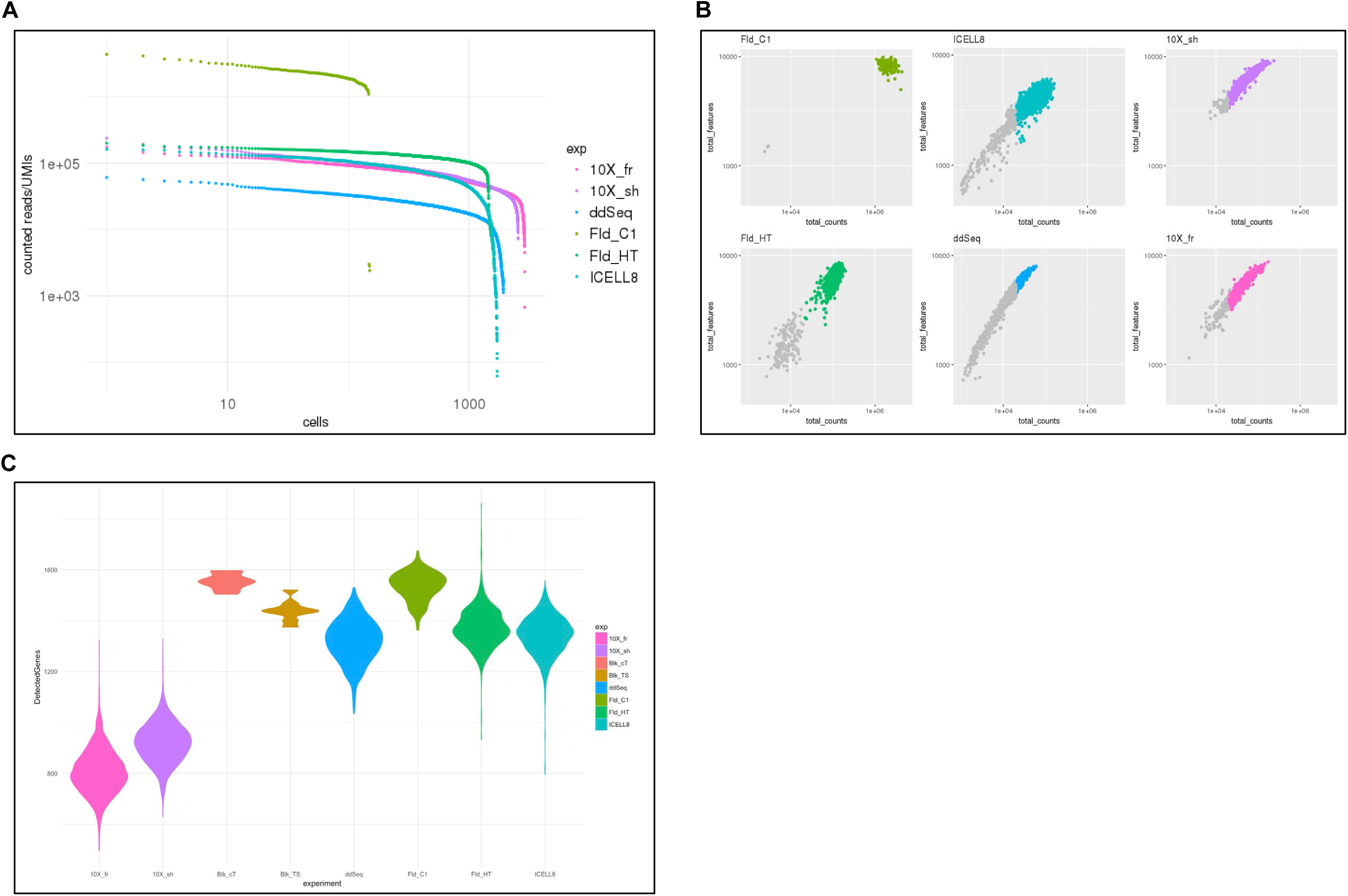
Sequencing saturation analysis. A: Cells were sorted by decreasing number of reads and the number of reads contributing to expression values is plotted. The Fluidigm C1 technology has more reads per cell but less cells than the other technologies. B: Relationship of the number of genes detected in a cell vs the number reads contributing to the expression values. C: Violin plots showing the number of genes detected when the sequencing depth is subsampled to 20’000 reads for each cell.

### Physical gene properties impact capture efficiency of scRNA-seq methods

Previous reports suggest that RNA-seq methods are biased with respect to gene length and GC content of the genes^11, 12, 13, 14, 15^, for example, genes with high GC content tend to be underrepresented. We investigated whether certain gene properties, (e.g. GC content and gene length) impact the efficiency with which the single-cell technologies capture genes, thereby imparting platform specific bias in these data. As shown in Figure 4 top panels, single cell platforms Fluidigm HT and ICELL8 show a lower capture efficiency for high GC content genes than of reference genes which we defined as genes with a GC content in the interval 0.5—0.55. Moreover, these data suggest that the 10X platform has a much lower bias for high GC content genes relative to the other technologies, more similar to bulk RNA-seq data. Interestingly, the ddSEQ platform has reduced capture efficiency for both high-GC and low-GC content genes, while Fluidigm and ICELL8 have higher capture efficiency for low-GC genes. We also visualized the relative capture efficiency for gene length bias (short/long gene). These data suggest that 3’-tagging technologies tend to over-represent short genes, with Fluidigm-HT having the smallest length bias and 10X having the strongest observed bias (Figure 4, bottom panels). While all single-cell technologies tend to under-represent long genes, Fluidigm platforms (Fluidigm C1 and Fluidigm HT) exhibit the least bias in that respect. Particularly interesting is that the Fluidigm HT platform performs in our experiment better across genes of varying length than Fluidigm C1. This is despite the fact that Fluidigm HT detects reads next to the 3’-end only (3’-end sequencing) while Fluidigm C1 detects genes along the entire gene body.

**Figure 4:**
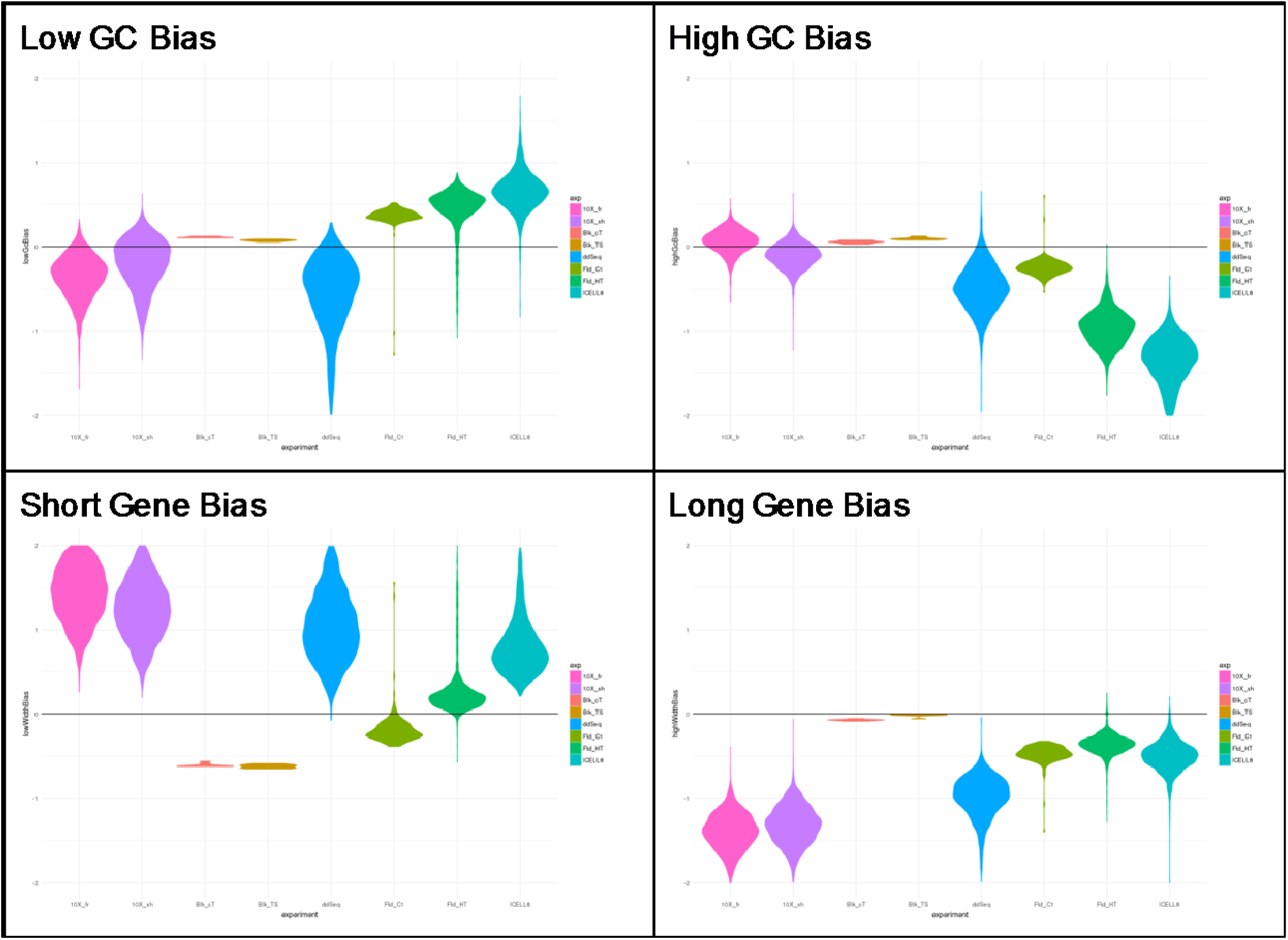
Detection bias of low-GC, high-GC, short and long genes. The violin plots show the bias present in the individual cells. For reference we added also the bulk protocols. Interestingly short genes are underrepresented in bulk RNA-seq and Fluidigm C1 but not in the other scRNA-seq technologies. Overall the bulk technologies show the smallest bias.

### All technology platforms have a high degree of specificity on protein coding regions of transcripts

A commonly utilized metric indicating the quality of RNA-seq data, is the percentage of data aligned to protein-coding regions of transcripts. The basic concept is that in poor quality RNA-seq data, more data aligns to regions outside of protein-coding regions, such as intergenic regions of the genome. To assess the quality of data produced by each scRNA-seq platform, we determined the proportion of data identified as various gene classes (Figure 5). All technologies exhibited good specificity with over 90% of the data assigned to protein-coding genes (Figure 5A). Figures 5B and 5C demonstrate the percentages of the data across platforms that are assigned to other gene types, such as long non-coding RNA (lncRNA) and small RNA species. Surprisingly, the 10X platform contained more antisense and miscRNA assigned data than the other platforms (Figure 5E-G). This observation correlates with the high capture efficiency of short protein coding genes of the 10X platform (Figure 4). These results suggest that while all technologies perform well with regard to detecting and quantifying protein-coding transcripts, some are better suited to identify poly-adenylated lncRNAs that may be of interest for specific experimental goals.

**Figure 5:**
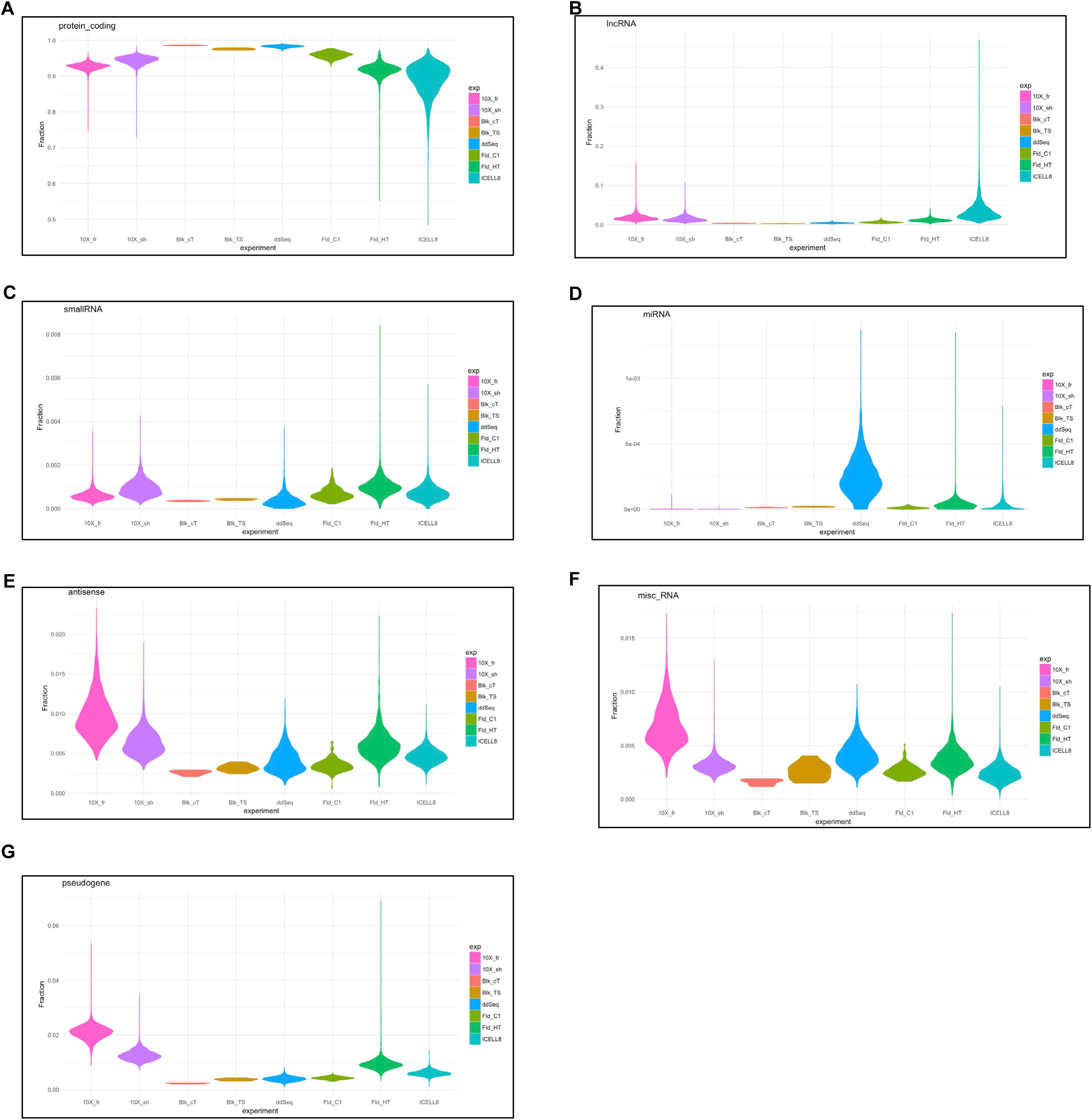
Distribution of gene types detected by the different technologies.

### Normalization and filtering of high-quality cells

In the subsequent sections we evaluated the cross-technology concordance of the gene expression matrices. In particular, we computed the correlation of the gene expression profiles between the different single-cell technologies and the bulk RNA protocols. Additionally, we evaluated the consistence of the expression ratios computed between the two treatment conditions TSA and DMSO. To this end we filtered and normalized the expression data using the scater package. Importantly, this analysis was done using the protein-coding genes only. For the concordance analysis we used the illumina TruSeq data as reference.

### Concordance of gene expression

To begin evaluating concordance across scRNA-seq platforms, we first determined the level of gene expression correlation within the DMSO control samples relative to the mean expression of the Illumina bulk TruSeq data. As expected, the three replicate DMSO samples processed with Illumina TruSeq data are highly concordant with their mean, correlation values close to 1.0 (Figure 6). In addition, the bulk samples generated with the Clontech low input protocols also show high correlation, around 0.96 (Figure 6). However, the correlation across the individual cells from each scRNA-seq platform with bulk data is considerably lower with values between 0.5 and 0.8 (Figure 6). Highest correlation is found for Fld_C1 and lowest correlation is found for ICELL8. 10X, ddSEQ and Fld_HT perform similarly with respect to this measure (Figure 6). Similar results are found when stratifying the capture rates of the individual technologies against the bulk RNA-seq expression (Figure 6). For any given expression level in bulk, Fld_C1 has the highest capture rate while ICELL8 has the lowest capture rate.

**Figure 6:**
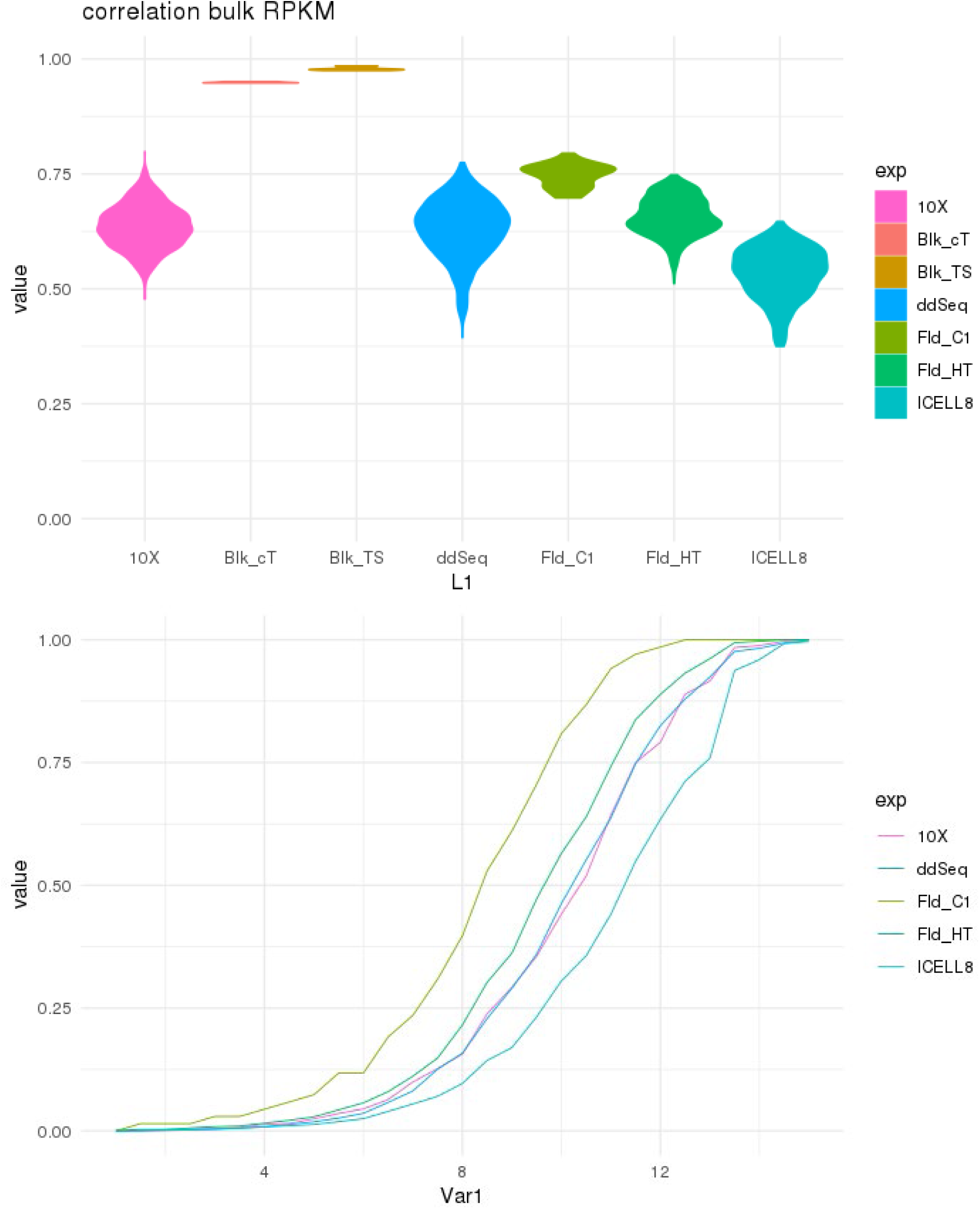
A) Correlation of single cell expression profiles with the bulk RNA-seq profile. B) Gene detection rate stratified by bulk RNA-seq expression. The plot shows the median detection rate in each strata. The x-axis shows the log2 read counts in the bulk RNA-seq data.

In order to assess how well expression differences between populations of cells can be quantified by the various single-cell technologies, we computed the expression difference between TSA and DMSO samples. Again, we used the Illumina bulk TruSeq data as the gold standard against which the other technologies are compared. Figure 7A shows that the two bulk RNA-seq technologies, TruSeq vs Clontech, do not only correlate well with respect to their absolute expression levels, however, both technologies are consistent with respect to the expression profile differences (Figure 7B). When looking at the single cell technologies, the correlation values of the expression changes are markedly lower (Figure 7B). Here we find that the expression differences of Fld_HT are most in-line with expression differences observed with the bulk RNA-seq methods (Figure 7B). As a noteworthy result, we find that for the Fld_C1 technology, the expression differences between TSA and DMSO treated samples have a low correlation with the remaining single-cell technologies as well as with the bulk RNA-seq samples.

**Figure 7:**
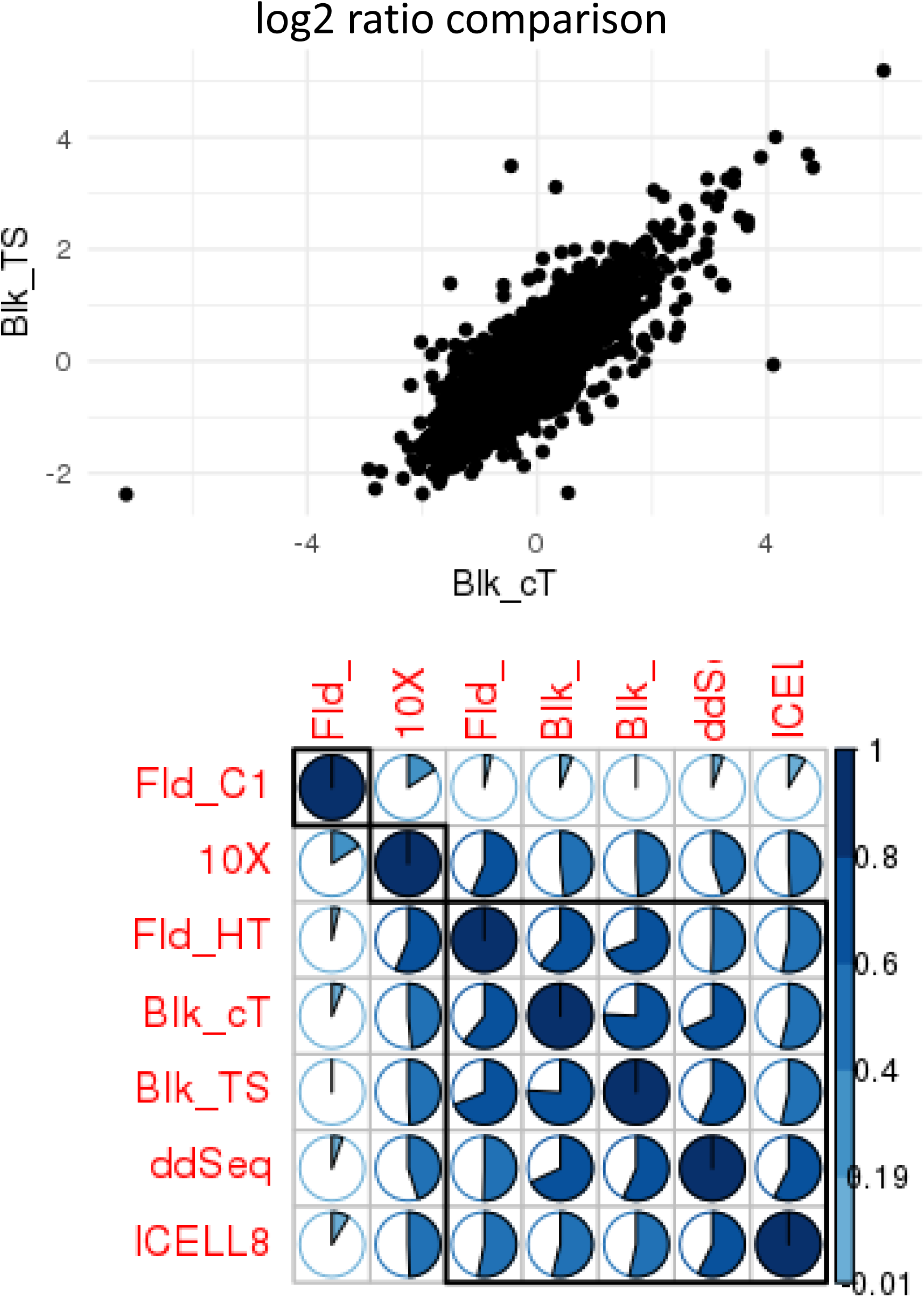
Correlation of expression ratios TSA vs DMSO. A) Scatter plot comparing the two bulk RNA-seq technologies. B) Correlation plot showing the pair-wise correlation values as pie charts

### Highly variable genes (HVGs) significantly overlap across platforms

A key feature of single-cell technologies is the ability to capture expression heterogeneity among cells. And in fact, for this type of analysis the most informative genes are those that are highly variable across cells. Following a recent comparative study on the effectiveness of existing methods^10^, we chose the method implemented in Seurat^17^ to identify highly variable genes whose expression values have variability that is higher than expected given their average expression. Figure 8 shows pairwise overlaps, in terms of their Jaccard coefficient, have been used to cluster the sets of HVG genes from different platforms. Not surprising, the highest overlap is found between ddSEQ and 10X, the two droplet-based platforms, which form a cluster. The two Fluidigm technologies also form a cluster (Figure 8). Only the ICELL8 technology shows only a very small overlap with the other methods (Figure 8). Taken together, these results support the cross-platform differences inherent to each scRNA-seq technology evaluate in this study. Importantly, this highlights the potential need to tightly control technology utilized for large multi-center studies to allow for more accurate data integration.

**Figure 8:**
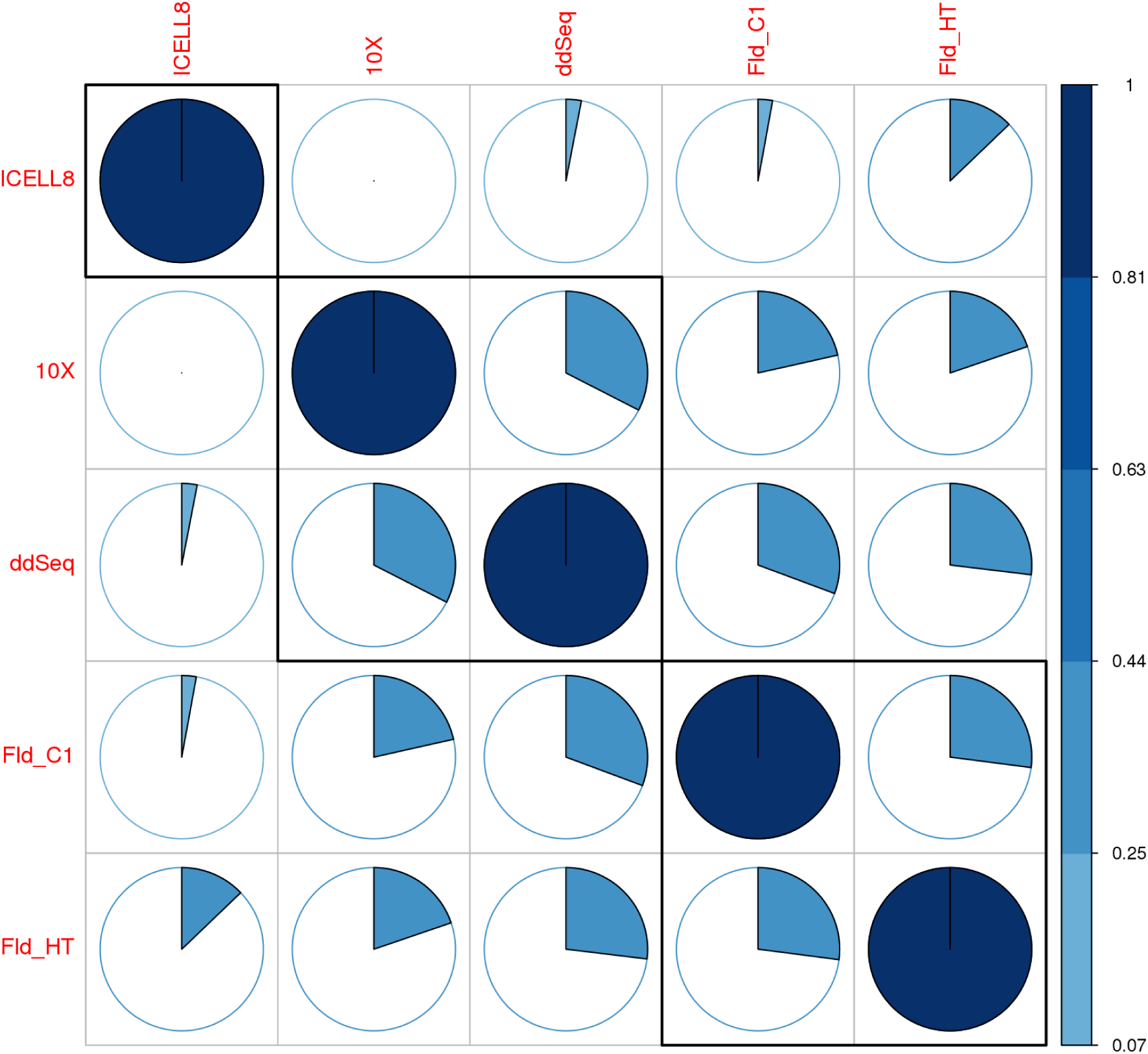
Comparing the highly variable genes across technologies. The plot shows the Jaccard Index of the pair-wise overlap of the highly variable genes.

### Availability

Data generated by all technologies have been uploaded to GEO (https://www.ncbi.nlm.nih.gov/geo/) and are available und the accession GSE142652.

## DISCUSSION

Single-cell RNA-seq technologies offer unprecedented resolution towards dissecting the cellular architecture of complex biological systems. The single-cell genomics landscape is rapidly evolving with various chemistries, platforms, and technologies currently in practice. Regardless of these differences, single-cell transcriptome profiling is a multi-stage process involving cell capture, efficient cell lysis, transcript capture, reverse transcription, cDNA preamplification, and final cDNA library construction suitable for high-throughput DNA sequencing. Understanding the intricacies and limitations of the various scRNA-seq modalities available is paramount to the success of the study. At time of this study, there were four commercial scRNA-seq platforms available for evaluation; Fluidigm C1, WaferGen/Takara iCell8, 10X Genomics Chromium Controller, Illumina/BioRad ddSEQ.

The goal of this comparative study was to demonstrate the level of reproducibility across several commercially available single-cell RNA sequencing (scRNA-seq) technologies using a well-characterized experimental model system for the purpose of highlighting the key characteristics of each platform. Our results suggest that while each scRNA-seq platform can provide detailed transcript information on the individual cell level, some biases exist that should be carefully considered during experimental design. Overall, the technologies do capture well the protein-coding genes within the limits of systematic detection biases; biases are present in terms of preferential sequencing high-GC and short genes. Despite of these biases the experiments do still provide valuable biological insight because low GC and long genes are still being detected. This bias does not hinder the comparison of individual gene expression levels across cells. Such a bias would however become relevant when directly comparing read counts expression across technologies or when relating read counts of two different genes to each other.

A key question is how compatible and comparable are the expression values obtained from individual cells with the traditional RNA-seq analysis of bulk tissue. In our analysis of SUM149PT cells, both approaches do correlate and give overall consistent results. We considered bulk RNA-seq as the gold standard of the single cell technologies and performed two well established protocols, the Illumina TruSeq protocol and the Clontech protocol. Both protocols gave highly consistent results and which supports the idea of using bulk RNA-seq as reference. When reporting correlation values with single cell protocols, we use the TruSeq protocol as reference and not the Clontech protocol because the latter shares some library preparation steps with the Fld_C1 protocol, which might favor the Fld_C1 protocol in the performance evaluation.

Our study involved two different treatments, TSA and DMSO, of the cells and permitted the computation of expression differences and the evaluation of their consistency. Overall the bulk RNA-seq expression differences between the two treatments DMSO and TSA are also reflected in the single cell data.

A key characteristic of single cell technologies is the throughput in terms of the number of cells and the associated cell success rate. The capacity of the microfluidic technology is lower than for droplet and plate-based technologies with the later technologies providing the possibilities to scale up the experiments. The higher capacity however comes with a compromise regarding the cell success rate. While the microfluidic chips reach up to 90% success rate in our study this is not true for the other systems. For them, we observe only a success rate that ranges between 50 to 80%. These numbers certainly depend on the cell types and cell viability. As a consequence that cannot be generalized to any cell type and any condition. An important lesson is that researchers should factor in failing cells when designing single cell studies and plan the number of cells entering the study accordingly.

Of equal importance is the number of genes detected per cell, which represents a characteristic complementary to the number of cells. Only when transcripts are detected with a high sensitivity in a cell the full potential of scRNA-seq unfolds. In terms of gene detection sensitivity, the microfluidics platforms perform very well. They have the highest number of detected genes. The other platforms that do measure many more cells do perform less well regarding the gene detection sensitivity per cell. We are aware that the detection sensitivity also depends on the sequencing depth, since transcripts may not only be lost at cDNA and subsequent library generation but also in the sequencing step that acts as a random sampling process. So, by putting more sequencing efforts into the experiment, gene detection can be improved until the limits imposed by the library generation are reached. The downside are the increasing costs and the inefficiencies caused by sequencing duplicated reads. Our general finding that the sequencing is not yet saturated is consistent with Ziegenhain et al.^7^ that shows an increase in gene detection sensitivity for sequencing depths up to approximately two million reads/cell for protocols doing full transcript sequencing.

Our summary of the single cell data demonstrates the capabilities of the technologies and sheds also light on the limitations. We abstain from a performance rating of the technologies since we have evaluated them only based on a single experiment using one cell type. Our analyses highlight the considerations to be taken into account when planning, executing and especially analyzing single cell data

## ACKNOWLEDGEMENTS

The authors would like to thank Dr. Martin Tenniswood, for generous donation of the SUM149PT cells. We would also like to acknowledge the generous donation of kits and reagents by 10xGenomics, BioRad, Fluidigm, Illumina and Wafergen Biosystems,

The authors declare no conflict of interest.

## REFERENCES

1. Karaayvaz, M. et al. Unravelling subclonal heterogeneity and aggressive disease states in TNBC through single-cell RNA-seq. Nat Commun 9, 3588 (2018).

2. Navin, N., et al. Tumour evolution inferred by single-cell sequencing. Nature, 472(7341), 90–94 (2011).

3. Sen, R. et al. Single-Cell RNA Sequencing of Glioblastoma Cells. Methods Mol Biol 1741, 151–170 (2018).

4. Saunders, A. et al. Molecular Diversity and Specializations among the Cells of the Adult Mouse Brain. Cell 174, 1015–1030 e1016 (2018).

5. Goldstein, L.D. et al. Massively parallel nanowell-based single-cell gene expression profiling. BMC Genomics 18, 519 (2017).

6. Baran-Gale, J., Chandra, T. & Kirschner, K. Experimental design for single-cell RNA sequencing. Brief Funct Genomics 17, 233–239 (2018).

7. Ziegenhain, C. et al. Comparative Analysis of Single-Cell RNA Sequencing Methods. Mol Cell 65, 631–643 e634 (2017).

8. Chatterjee, N., Wang, W.L., Conklin, T., Chittur, S. & Tenniswood, M. Histone deacetylase inhibitors modulate miRNA and mRNA expression, block metaphase, and induce apoptosis in inflammatory breast cancer cells. Cancer Biol Ther 14, 658–671 (2013).

9. Hicks, S.C., Townes, F.W., Teng, M. & Irizarry, R.A. Missing data and technical variability in single-cell RNA-sequencing experiments. Biostatistics 19, 562–578 (2018).

10. Hwang, B., Lee, J.H. & Bang, D. Single-cell RNA sequencing technologies and bioinformatics pipelines. Exp Mol Med 50, 96 (2018).

11. Zheng, W., Chung, L.M. & Zhao, H. Bias detection and correction in RNA-sequencing data. BMC Bioinformatics 12, 290 (2011).

12. Tuerk, A., Wiktorin, G. & Guler, S. Mixture models reveal multiple positional bias types in RNA-seq data and lead to accurate transcript concentration estimates. PLoS Comput Biol 13, e1005515 (2017).

13. Young, M.D., Wakefield, M.J., Smyth, G.K. & Oshlack, A. Gene ontology analysis for RNA-seq: accounting for selection bias. Genome Biol 11, R14 (2010).

14. Roberts, A., Trapnell, C., Donaghey, J., Rinn, J.L. & Pachter, L. Improving RNA-seq expression estimates by correcting for fragment bias. Genome Biol 12, R22 (2011).

15. Oshlack, A. & Wakefield, M.J. Transcript length bias in RNA-seq data confounds systems biology. Biol Direct 4, 14 (2009).

16. Soneson, C. & Robinson, M.D. Bias, robustness and scalability in single-cell differential expression analysis. Nat Methods 15, 255–261 (2018).

17. Satija, R., Farrell, J.A., Gennert, D., Schier, A.F. & Regev, A. Spatial reconstruction of single-cell gene expression data. Nat Biotechnol 33, 495–502 (2015).

